# Residue-Level Allostery Propagates Through the Effective Coarse-Grained Hessian

**DOI:** 10.1101/822882

**Authors:** Peter T. Lake, Russell B. Davidson, Heidi Klem, Glen M. Hocky, Martin McCullagh

## Abstract

The long-ranged coupling between residues that gives rise to allostery in a protein is built up from short-ranged physical interactions. Computational tools used to predict this coupling and its functional relevance have relied on the application of graph theoretical metrics to residue-level correlations measured from all-atom molecular dynamics (aaMD) simulations. The short-ranged interactions that yield these long-ranged residue-level correlations are quantified by the effective coarse-grained Hessian. Here we compute an effective harmonic coarse-grained Hessian from aaMD simulations of a benchmark allosteric protein, IGPS, and demonstrate the improved locality of this graph Laplacian over two other connectivity matrices. Additionally, two centrality metrics are developed that indicate the direct and indirect importance of each residue at producing the covariance between the effector binding pocket and the active site. The residue importance indicated by these two metrics is corroborated by previous mutagenesis experiments and leads to unique functional insights; in contrast to previous computational analyses, our results suggest that fP76-hK181 is the most important contact for conveying direct allosteric paths across the HisF-HisH interface. The connectivity around fD98 is found to be important at affecting allostery through indirect means.

## 1 Introduction

Allostery refers to the long-range functional coupling of sites in a macromolecule through networks of short-ranged interactions. This phenomenon can be a pivotal component of a protein’s function ^1^ as demonstrated in GPCR signaling, ^2^ coupling of ATP binding and hydrolysis to mechanical work in motor proteins ^3–6^ and activation of oxygen binding in hemoglobin. ^7–12^ Many of these processes are initiated by the binding of an effector molecule that modulates activity at a distal active site. Allosteric pockets have become an increasingly important target in the field of drug development due to possible improvements in selectivity over orthosteric sites. ^13–17^ Intriguingly, allostery can be incorporated into an enzyme’s function even on the short time scales of directed evolution studies to produce a desired increase in activity. ^18^ Therefore, it is highly desirable to be able to identify and predict allostery, as well as the interactions (“allosteric paths”) involved in these processes.

There are many well established experimental techniques for characterizing allostery, including activity based assays to investigate non-Michaelis-Menten kinetic behavior, ^19^ H/D mass spectrometry, ^20,21^ as well as structural based approaches such as X-ray crystallography, ^22^ cryoEM, ^23^ and NMR. ^24–32^ Structural techniques can be used to identify residues that interact directly with or are structurally perturbed by the effector molecule by comparing *apo* and effector bound states of a protein. Identifying allosteric paths is significantly more challenging. While paths have been identified using experimental techniques in well-studied proteins such as hemoglobin, ^10,33–35^ the combination of NMR spectroscopy and computational techniques have allowed for the most robust description of allosteric paths. ^27,30,31,36,37^

Computational techniques used to investigate allostery rely on graph theoretical approaches to identify important residues or connections that convey information from the effector binding pocket to the active site. ^29,38–40^ A weighted graph is constructed by defining nodes (residues) and edges (bonds) that connect nodes. These edge weights populate a pairwise adjacency matrix, ***A***, that has finite values between connected residues. The closely related graph Laplacian, ***L*** = ***D*** − ***A*** where ***D*** is a diagonal matrix with elements *D*_*ii*_ = Σ_*j*_*A*_*ij*_, can be readily constructed from the adjacency matrix. The definition of nodes and edge weights are what distinguish different computational techniques targeted at understanding allostery.

Two computational techniques have been used to generate these graphs: bioinformatics and molecular dynamics. Sequence-based bioinformatic methods, working under the assumption that allosteric pathways are functionally important and thus evolutionarily conserved, use residue pairwise sequence co-evolution to define a graph. ^29,35,41^ These methods do not explicitly take into account the 3D structure of a given protein as this information should be implicitly captured in sequence covariance. Additionally, structural databases can be used to build pairwise, knowledge-based potentials that describe the interactions observed in e.g. a protein’s crystal structure. ^42^ None of these models, however, directly account for the ensemble nature of protein structure and the associated allosteric behavior. ^11^

All-atom molecular dynamics (aaMD) is a well-established method to sample the configurational ensemble of a protein. ^43–48^ Analyses of such simulations have been used to identify allosteric importance based on residue interaction networks ^29,49–51^ or positional covariance-based metrics. ^36,37,51–53^ Both of these models are used to describe the average residue-residue couplings from the ensemble of protein configurations observed. Interestingly, these methods lead to dramatically different graphs: the residue interaction networks will be localized in space while the positional covariance will be delocalized. Thus, while aaMD provides an appealing measure of protein configurational space, the appropriate graph Laplacian to describe residue-level correlations is not well understood.

Given a graph, centrality metrics can be used to identify important nodes or edges in the graph. A variety of edge and node centrality metrics have been used to study allostery including closeness, betweenness, degree and eigenvector. ^29,37,54^ These metrics are well defined for a given graph but their importance for allostery is unclear. Degree centrality, for example, provides a measure of the connectivity of a given node. A highly connected node is deemed to be important in a graph but how important is it for allostery between two given pockets? The degree centrality metric cannot directly report on allostery because it does not directly account for it. Similar arguments can be made for the other centrality metrics. Thus, it remains an open question in the field as to what centrality metric to apply to a given graph to better understand the allostery in a system.

In the next section, we demonstrate that the effective coarse-grained Hessian is the appropriate graph Laplacian to quantify residue-level allostery from aaMD simulations. Additionally, we develop two node centrality metrics that quantify importance by measuring how nodes in the graph convey or affect the covariance between a set of source and sink nodes. Finally, we apply this method to simulations of the imidazole glycerol phosphate synthase (IGPS) heterodimer, which is a well established benchmark allosteric system. *Our definition of the graph Laplacian and physically motivated centrality metrics provide new insight into the physics underlying allostery, a key functional component of most proteins.*

## 2 Theoretical Framework

Structural allostery can be described as the positional change at one site due to the application of an external force at a different site. A linear response of a system to an external force, ***f***_***ex***_, can be written as

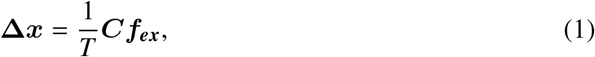

where **Δ*x*** is the change in the equilibrium position due to the perturbation, ***C*** is the matrix of covariance of particle positions, and *T* is temperature in units of energy. ^55^ Equation 1 demonstrates the importance of positional covariance to predicting allostery, but does not indicate anything about the short-ranged physical interactions leading to this behavior.

Within a graph theoretical framework, the short-ranged interactions that lead to long-ranged correlations are what should populate the graph Laplacian of the system. The Hessian is the appropriate graph Laplacian for molecular systems because (1) Hessian elements are only finite for short-ranged physical interactions, (2) the Hessian is, by construction, a symmetric positive semi-definite matrix as all graph Laplacians are, and (3) the covariance is related to the generalized inverse of the Hessian. The final point is most well-known in elastic network models (ENMs) for which the covariance and Hessian are related by the Moore-Penrose pseu-doinverse 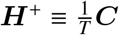. This relationship is consistent with the Gaussian Markov random field literature in which the graph Laplacian and the covariance are related by pseudoinverses. ^56^ In the context of allostery in proteins, the covariance is typically measured at a residue-level even if the underlying simulations have atomic resolution. Thus, the Hessian of interest is the second derivative of an effective coarse-grained Hamiltonian. Determining the effective Hessian from a measured covariance is a non-trivial problem due to the difficulties in converging a measured covariance. ^57^ Here we use a previously described coarse-graining procedure to generate an effective harmonic Hessian based on a measured covariance.

### 2.1 The Effective Harmonic Hessian from All-Atom Molecular Dynamics Simulations

Explicit solvent aaMD simulations can provide a detailed and accurate structural ensemble picture of proteins under physiological conditions. ^43–48^ These simulations have been used to generate 3*N* × 3*N* covariance matrices that have been used to assess *N*−residue level structural allostery. Motivated by the idea that the structural ensemble in a single free energy well from aaMD is well represented by a harmonic system, ^58–62^ effective harmonic Hamiltonians have been fit to mapped all-atom data. ^63–65^ In this work, we construct an effective harmonic Hessian using a slightly modified heterogeneous ENM (hENM) procedure ^64^ as described in the Supplementary Information (SI) Computational Methods section.

The result of the hENM procedure is a *N* × *N* force constant matrix, ***k***, that is optimized to reproduce pairwise particle variances. The 3 × 3 tensor element of the 3*N* × 3*N* Hessian is defined as

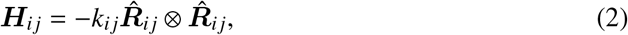

Where 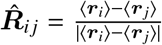 and 〈***r***_***i***_〉 is the average position of node *i*. The 3*N* × 3*N* covariance can then be reconstructed as ***C*** = *T* ***H***^+^.

Isotropic ENMs have been used to define a residue-level coarse-grained Hessian and study the contributions of the normal modes to allostery for a variety of systems. ^66^ These ENM models qualitatively capture low frequency motions of proteins, yet the lack of residue specificity in the model yields low fidelity with all-atom models in the mid to high frequency range. ^67^ hENM is an anisotropic ENM that is designed to capture higher fidelity with all-atom normal modes across the spectrum ^68^ as can be observed by the agreement between hENM covariance and aaMD measured covariance for IGPS depicted in Figure S1. By using an hENM-produced Hessian to study allostery, we are connecting the fields of protein allostery and coarse-grained potentials.

### 2.2 Allosteric Paths in the Hessian

To investigate allosteric paths, we start by dictating that the sum over these paths yields the covariance between a selected pair of source and sink residues. This is motivated by the idea that the covariance is the physical observable that dictates the linear response between two residues. Employing a property from graph theory that, given a weighted adjacency matrix, ***A***, (***A***^***ℓ***^)_*ij*_ is the weighted sum of all walks of length *ℓ* between nodes *i* and *j*, we can express the covariance in the following way,

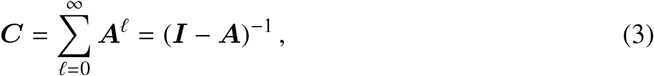

where *I* is the identity matrix and the last equality arises from identifying the infinite sum as an example of the Neumann series.

If we consider ***C*** to be strictly invertible and map the infinite number of walks to a finite set of paths, *P*_*ij*_, it can be shown that

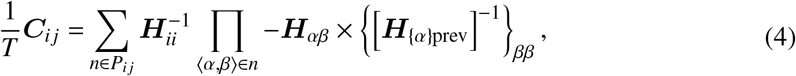

where 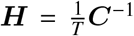, 〈*α, β*〉 is an edge in the path and subscript {α}prev denotes the principle submatrix of ***H*** obtained by removing all nodes previously visited in the path. The 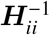 and [***H***_{α}prev_]^−1^ terms in Equation 4 arise from mapping walks to paths; traversing all loops from *α* to itself results in terms related to the conditional variance of *α*. The only terms that couple two nodes together in Equation 4 are ***H***_*αβ*_ which are the terms in the Hessian for a harmonic system.

Paths are usually referenced in terms of a length e as opposed to a weight. This distinction can be made by considering a path length as the negative of a sum of the adjacency matrix on a logarithmic scale as opposed to the weight, which is a product of elements in the adjacency matrix. In this way, the paths with the largest weights have the shortest lengths. We study these paths by sampling them in a Monte Carlo scheme; even though the paths observed are not necessarily all the shortest paths within some number, the algorithm is much more efficient than the typically used Floyd-Warshall algorithm. ^69,70^ This method of sampling paths also leads us to consider these paths as part of an ensemble, all of which can contribute. The Monte Carlo sampling is also readily extended to account for path lengths which are not strictly a sum of pairwise interactions. The procedure is detailed in the SI Theory section.

In the case of aaMD simulations of proteins, elements in the covariance and Hessian matrices are 3 × 3 tensors. Additionally, both matrices are singular and thus not strictly invertible. A similar but more complicated derivation can be followed to determine paths that yield the covariance in a singular covariance matrix. The most crucial idea obtained from performing the full derivation is that fully connected paths, those with finite Hessian values connecting a given source and sink and quantified in Equation 4, are not the only contributions to the covariance. Additional terms, that can be described as broken paths, contribute to the covariance due to a coupling through the null-space of the singular matrix. The relevant outcome of this for the current work is that residues that contribute to direct paths are not the only residues that affect the covariance between a source and sink. This motivates the need for additional importance metrics to study allosteric interactions in a given graph.

### 2.3 Hessian Derivative as a Centrality Metric

The effective harmonic Hessian lends itself to another centrality metric. One can consider how the covariance between a given set of sources and sinks is affected by changing a single spring constant. This leads to an edge-based centrality metric we call the derivative metric, defined by

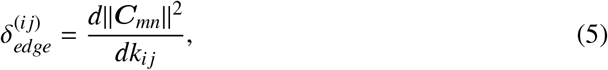

where ||***C***_*mn*_||^2^ is the squared Frobenius norm of the covariance tensor between source *m* and sink *n*. We map this metric down to a node-based centrality metric by

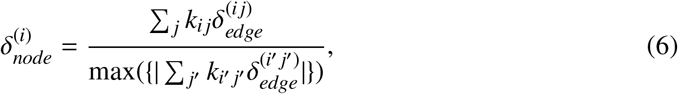

where the denominator represents a scaling factor to yield: 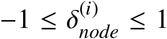. Observed values of 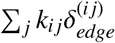 from our simulations ranged from −5.57 × 10^−3^Å^4^ to 2.20 × 10^−4^Å^4^ thus a scaling factor of 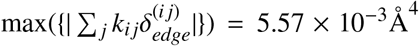 was used. The multiplication of the edge metric by the respective spring constant in Equation 6 to yield a node metric is motivated by considering a fractional change of all the spring constants connecting a given node.

### 2.4 Error Analysis

There are two major sources of error, other than force field concerns, in estimating the two centrality metrics mentioned above. The first of these is sampling of the residue-level covariance from aaMD simulations. We estimate this error by performing our analyses on chunked segments of trajectory (four 250 ns chunks) and computing the standard error from these four values. The averages reported are determined from applying our analyses on the covariance from the amalgamated trajectory. The second source of error is the inability to represent the simulation data with a coarse-grained harmonic Hamiltonian. This error is more difficult to quantify and is left for future study.

## 3 Results and Discussion

### Model protein - IGPS

Imidazole glycerol phosphate synthase (IGPS) is an enzyme that functions in both the purine and histidine biosynthesis pathways of plants, fungi, archaea, and bacteria. In *Thermatoga maritima*, IGPS is a heterodimeric protein complex of HisH and HisF proteins (depicted in white and orange respectively in Figure 1). HisH catalyzes the hydrolysis of glutamine into ammonia and glutamate. Nascent ammonia is then shuttled across the HisF–HisH interface and through the (β/α)_8_ barrel of HisF (an approximate distance of 25 Å). At the HisF cyclase active site, the ammonia reacts with N’-[(5’-phosphoribulosyl)formimino]-5-aminoimidazole-4-carboxamide ribonucleotide (PRFAR) to form imidazole glycerol phosphate (IGP) and 5-aminoimidazole-4-carboxamide (AICAR). These two reactions are strongly coupled with a 1:1 stoichiometry, despite the large distance that separates the two active sites. ^71,72^ In addition to the concerted mechanism between HisF and HisH, IGPS is classified as a V-type allosteric enzyme, such that the rate of the glutaminase reaction is critically dependent on the presence of the PRFAR ligand. Experimental assays have quantified this strong allosteric activation to be an approximate 4,900-fold increase in activity relative to basal levels. ^73^ Allosteric activation is also observed in the presence of the cyclization products, IGP and AICAR, but at reduced magnitudes. ^73,74^

**Figure 1:**
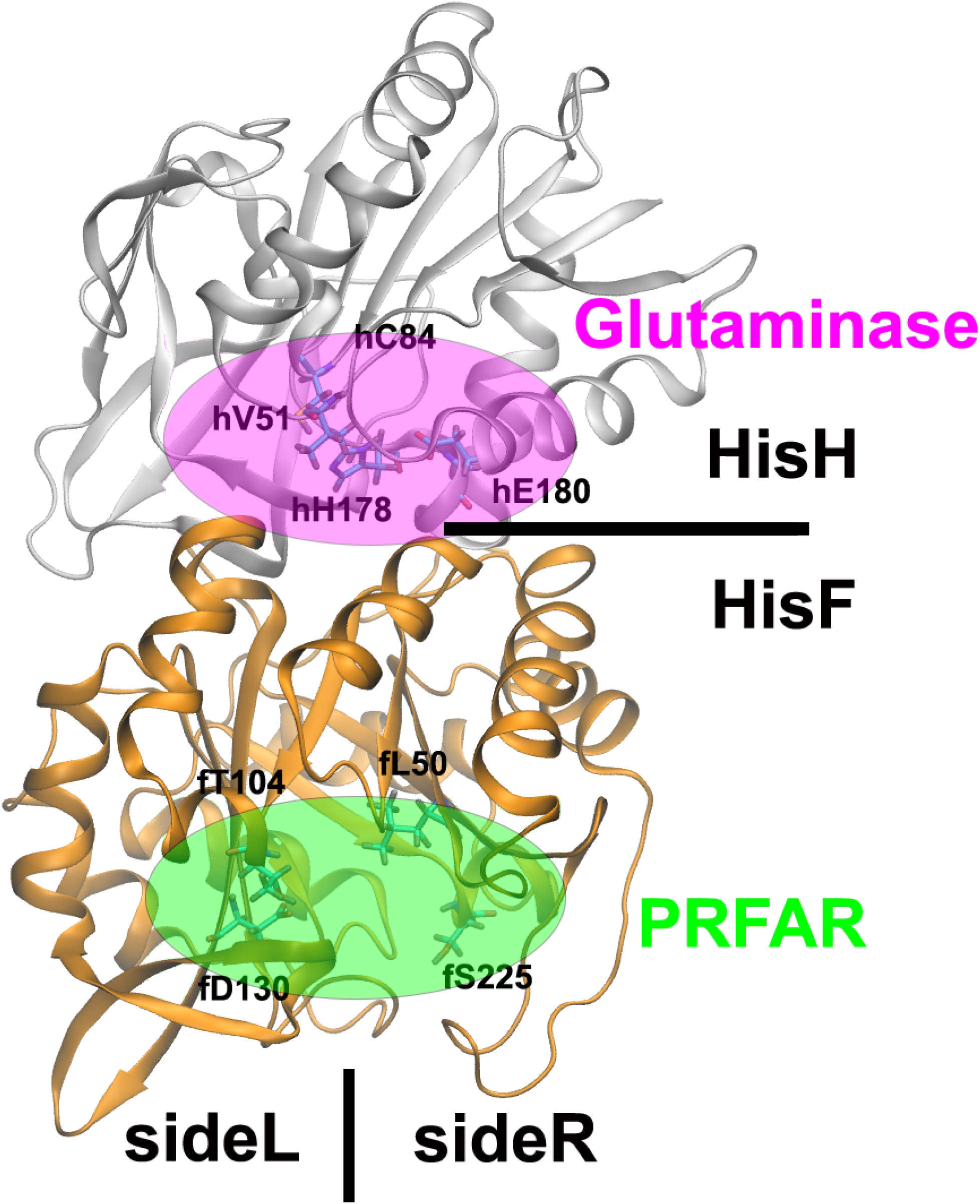
Structural depiction of the IGPS heterodimer. The protein is composed of HisH and HisF monomers. The PRFAR ligand binds in the pocket indicated by the green oval and source residues fL50, fT104, fD130 and fS225. The glutaminase active site, labeled in pink, is located in the HisH monomer near the interface. We chose four sink residues hV51, hC84, hH178 and hE180 to identify this pocket.

The allostery of the IGPS enzyme has been studied for nearly two decades using a broad range of experimental and theoretical methodologies. Mutational studies, coupled with biochemical assays and NMR experiments, have highlighted residues that are pivotal for the functionality of the protein complex as well as dynamic effects induced by the HisF ligand. ^30,31,73–81^ Additionally, MD simulations and graph-theoretic analyses have been applied to the IGPS system to study the allosteric effect of PRFAR and the allosteric paths that couple the two distal active sites. ^30,36,37,51,52,76,82–85^ For this work, we have performed aaMD simulations of *apo* IGPS as well as one mutant variant, totalling 1µs of trajectory for each system. Subsequently, an effective harmonic Hessian has been constructed and analyzed for both systems (SI Computational Methods).

### 3.1 Adjacency Matrix Comparison

Adjacency matrices based on the covariance and mutual information (MI) have been used to study allostery in IGPS. ^51,52,84^ The theoretical framework outlined above demonstrates the importance of a third adjacency matrix, one based on the Hessian. In order to compare and contrast between these three, we consider absolute values of normalized adjacency matrices. These matrices are limited to values between 0 and 1, with values near one indicating strong connectivity and values near zero indicating weak connections. Residue-level normalized adjacency matrices based on our 1µs MD simulation of *apo* IGPS are shown in Figure 2.

**Figure 2:**
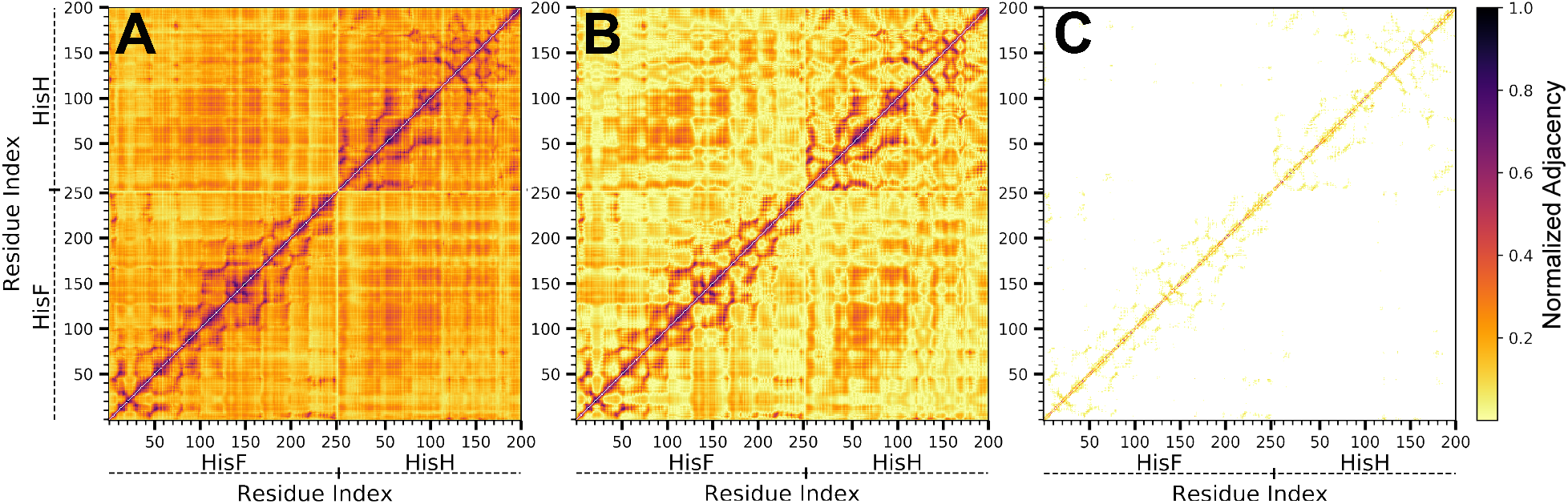
Normalized adjacency matrices computed from 4 × 250 ns all-atom molecular dynamics simulations of IGPS protein in its *apo* form. A) The linear mutual information adjacency, rMI,^86^ B) the Pearson correlation adjacency and C) the effective harmonic Hessian.

Mutual information (MI) is a measure of correlation between two distributions based on the difference between marginal and joint Shannon entropies. The numeric values of this quantity range from zero to infinity, but can be “normalized” by computing 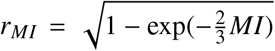 where this manipulation leads to pairs with large MI having *r*_*MI*_ ≈ 1 and uncoupled pairs having *r*_*MI*_ ≈ 0. We note, however, that neither the MI nor the *r*_*MI*_ matrices are positive semidefinite, which differentiates it from the two other matrices discussed here. The motivation for using this “adjacency” matrix over, for example, the normalized 1D covariance matrix, is that it accounts for correlation along perpendicular degrees of freedom. ^86^ The residue-level *r*_*MI*_ matrix, computed from a linear MI, for our *apo* IGPS simulations is depicted in Figure 2A. This plot indicates strong correlations along the primary sequence as indicated by dark colors in the near off-diagonal elements of the matrix. Further off-diagonal dark coloring indicates secondary structural elements that also convey strong correlation. The break between the HisF and HisH proteins is also evident as there are clear dividing lines at residue 253. Despite stronger near diagonal correlation, the *r*_*MI*_ adjacency matrix demonstrates long-range contacts even in the HisF–HisH coupling sub-matrix in the bottom-right or top-left of the matrix.

An alternative to *r*_*MI*_ is the Pearson correlation matrix. This adjacency matrix is derived from the covariance matrix, which is positive semi-definite. In a 3D system of *N* particles, the covariance matrix is a 3*N* × 3*N* matrix or, equivalently, an *N* × *N* matrix of 3 × 3 tensor elements. A typical manipulation is to reduce the 3*N* × 3*N* matrix to an *N* × *N* matrix by taking the trace of each tensor element. While this manipulation still yields a positive semi-definite matrix, we do note that it ignores couplings in orthogonal degrees of freedom upon mapping the system to a 1D system. ^86^ The resulting *N* × *N* matrix can be further manipulated to the Pearson correlation matrix with values between −1 and 1 by dividing by the square root of the product of diagonal elements of the corresponding row and column. Taking the absolute value of the Pearson correlation for the *apo* IGPS system yields the normalized adjacency matrix depicted in Figure 2B. While there are quantitative differences observed between the *r*_*MI*_ and Pearson correlation, the qualitative behavior remains the same: there are strong contacts along the primary sequence, but there are finite contacts for almost all elements of the matrix.

The third adjacency matrix we consider is based on the effective harmonic Hessian. We compute this from the 3*N* × 3*N* covariance matrix and the average structure from a simulation following the modified hENM procedure as discussed in the Theoretical Framework section and further elaborated on in the SI. The Hessian is a symmetric positive semi-definite matrix by construction, and the normalized adjacency matrix can be constructed in a manner analogous to what is done for the Pearson correlation. The Hessian-based normalized adjacency matrix (Figure 2C) shows strong primary sequence connectivity in agreement with the Pearson correlation and *r*_*MI*_ adjacency matrices. Secondary structure connectivity is observed in the near off-diagonal of Figure 2C in agreement with the strongly correlated off-diagonal regions of Figure 2A and B. These connections are distinctly weaker in the Hessian-based adjacency, however, and the farther off-diagonal elements are zero indicating little long-range connectivity in the graph. This adjacency matrix is consistent with the idea of short-ranged physical interactions that yield long-range correlation.

### 3.2 Direct Paths Comparison

Paths that convey the covariance between a set of sources and sinks are attractive physical interpretations of the interactions that couple distal sites in allostery. Direct paths, ones with complete connectivity in the adjacency matrix, are an incomplete picture of the covariance between any given source and sink, but are readily sampled. We use direct paths here to compare and contrast adjacency matrices. In this work, we sample paths between four source residues (fL50, fT104, fD130, fS225) that span the effector-binding pocket (green oval in Figure 1) and four sink residues (hV51, hC84, hH178, hE180) in the glutaminase active site (pink oval in Figure 1) of IGPS that have been previously identified as important. ^36,87,88^ The ensemble of paths from all 16 source-sink pairs are reweighted together to provide a comprehensive picture of the coupling between the two active sties. Path sampling is performed in a Monte-Carlo scheme as described in the Theoretical Framework section and SI.

Paths between sources and sinks in the Hessian-based adjacency matrix are longer and less degenerate than those observed in the Pearson or *r*_*MI*_ matrices. Figure 3A depicts the degeneracy of paths as a function of path length, *ℓ*, for the Pearson correlation, *r*_*MI*_ and Hessian paths. The paths in the Pearson correlation and *r*_*MI*_ adjacency matrices behave similarly: paths of extremely short length are found and the degeneracy grows rapidly with path length. This can be understood by observing finite values in all elements of these two matrices. The paths on the Hessian are longer and the degeneracy of paths grows much less rapidly than either the Pearson or *r*_*MI*_ paths. Again, this can be understood by the smaller number of connections in the adjacency matrix leading to both longer paths as well as more unique, short path lengths. These results demonstrate that the paths on the Hessian use a small number of short-ranged interactions to produce long-range correlation.

**Figure 3:**
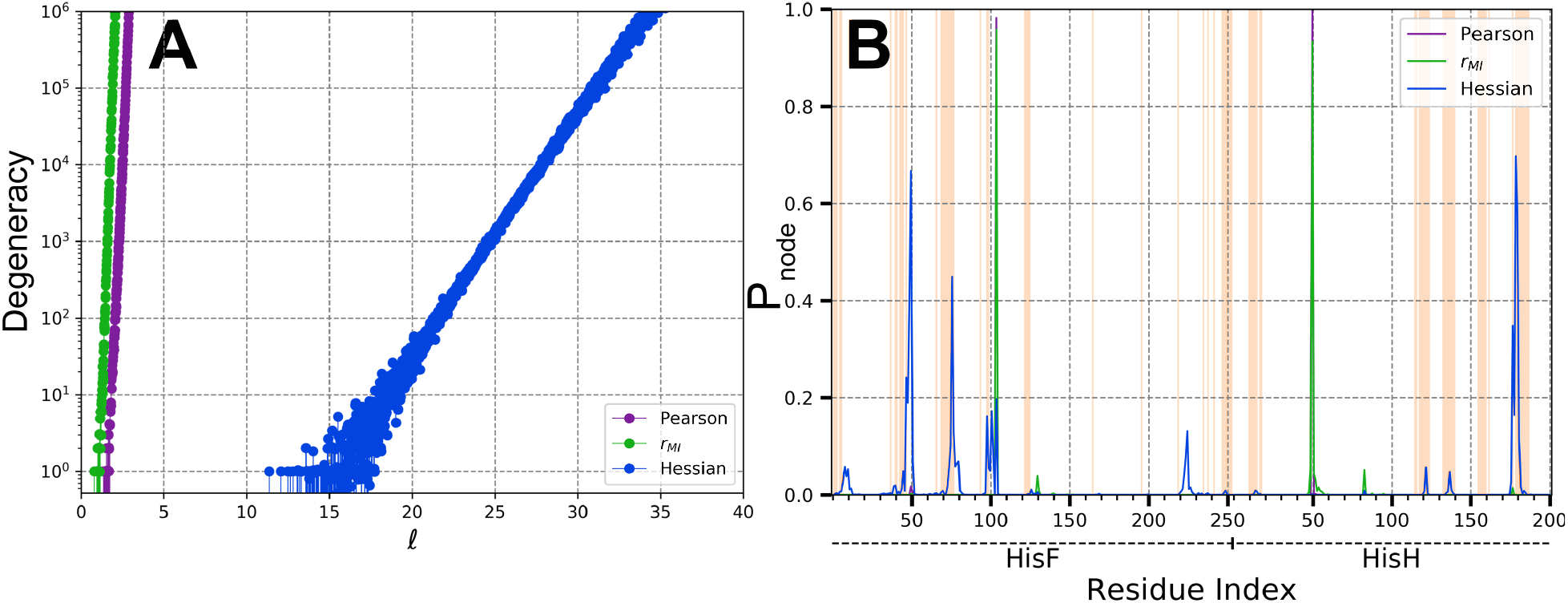
Comparison of paths and resulting centralities for different adjacency matrices for the IGPS protein. A) The path length degeneracy as a function of path length, *ℓ*. B) The probability of observing a given node in a sampled path, *P*_*node*_. Residues that have connections across the HisH-HisF interface are colored by vertical orange lines.

Paths in the Hessian sample different nodes than paths in the other two adjacency matrices. To quantify this, we define a centrality metric, *P*_*node*_, as the probability of observing each residue in the sampled paths. If applied to the Hessian, this metric represents a nodes importance at conveying the covariance between sources and sinks as indicated by Equation 4. We plot this metric as a function of residue number in Figure 3B for each adjacency matrix. The *r*_*MI*_ and Pearson adjacency matrices both have high probability of observing the source residue fT104 (*P*_*node*_ > 0.9) and sink residue hV51 (*P*_*node*_ > 0.9) in the paths but little probability of observing any other residue. This is due to the large covariance observed between residues fT104 and hV51 yielding large *r*_*MI*_ and Pearson adjacency values and thus direct path between the two; the two residues are measured to be ∼ 25Å apart. The Hessian-based adjacency matrix emphasizes the importance of source residues fL50 and fT104 and sink residues hE180 and hH178. Unlike paths from *r*_*MI*_ and Pearson adjacency matrices, paths in the Hessian also sample non-source and non-sink residues with high probability including hK181 and fP76. We thus conclude that the paths found in the Hessian yield much more physically relevant information that paths in the other two adjacency matrices.

The Hessian-based paths go through known important regions of IGPS. Figure 4A depicts the 100 shortest paths in IGPS, highlighting the localization of residues sampled in paths to sideR of HisF a region known to be important in the allosteric regulation of the enzyme. ^36,37^ The 100 shortest paths mainly propagate from fL50 (*P*_*node*_ = 0.68(2)) and fT104 (*P*_*node*_ = 0.18(3)) to hH178 (*P*_*node*_ = 0.33(5)) and hE180 (*P*_*node*_ = 0.71(5)). The paths from fL50 initially propagate through the backbone of f*β*2 to fF49 (*P*_*node*_ = 0.40(3)) and, to a lesser extent, fV48 (*P*_*node*_ = 0.19(3)). The paths jump from f*β*2 to f*β*3 through the hydrophobic pocket of fF49, fV48 and fF77 (*P*_*node*_ = 0.12(2)). The paths then propagate along the primary sequence of f*β*3 to fP76 (*P*_*node*_ = 0.43(4)) and then jump the HisF-HisH interface to hK181 (*P*_*node*_ = 0.59(6)) and end up at sink residues hH178 or hE180. We note that VanWart *et al.* found a similar path using a Cα mapped coarse-graining and a distance throttled Pearson correlation adjacency matrix. ^85^ Their path was disregarded in favor of an alternative path in part due to the presence of fT78, a poorly conserved residue, that does not show-up with large probability in our paths (*P*_*node*_ = 0.04(2)). Paths emanating from fT104 propagate through the backbone f*β*4 (fI102, fS101, fV100 etc) and either hop the HisF-HisH interface at fD98 (*P*_*node*_ = 0.16(6)) or jump to fP76 and then through the fP76-hK181 contact. Thus, we find that allosteric paths in IGPS are mainly propagated by sideR of the *β*-barrel of HisF before crossing the HisF-HisH interface mainly through the fP76-hK181 contact.

**Figure 4:**
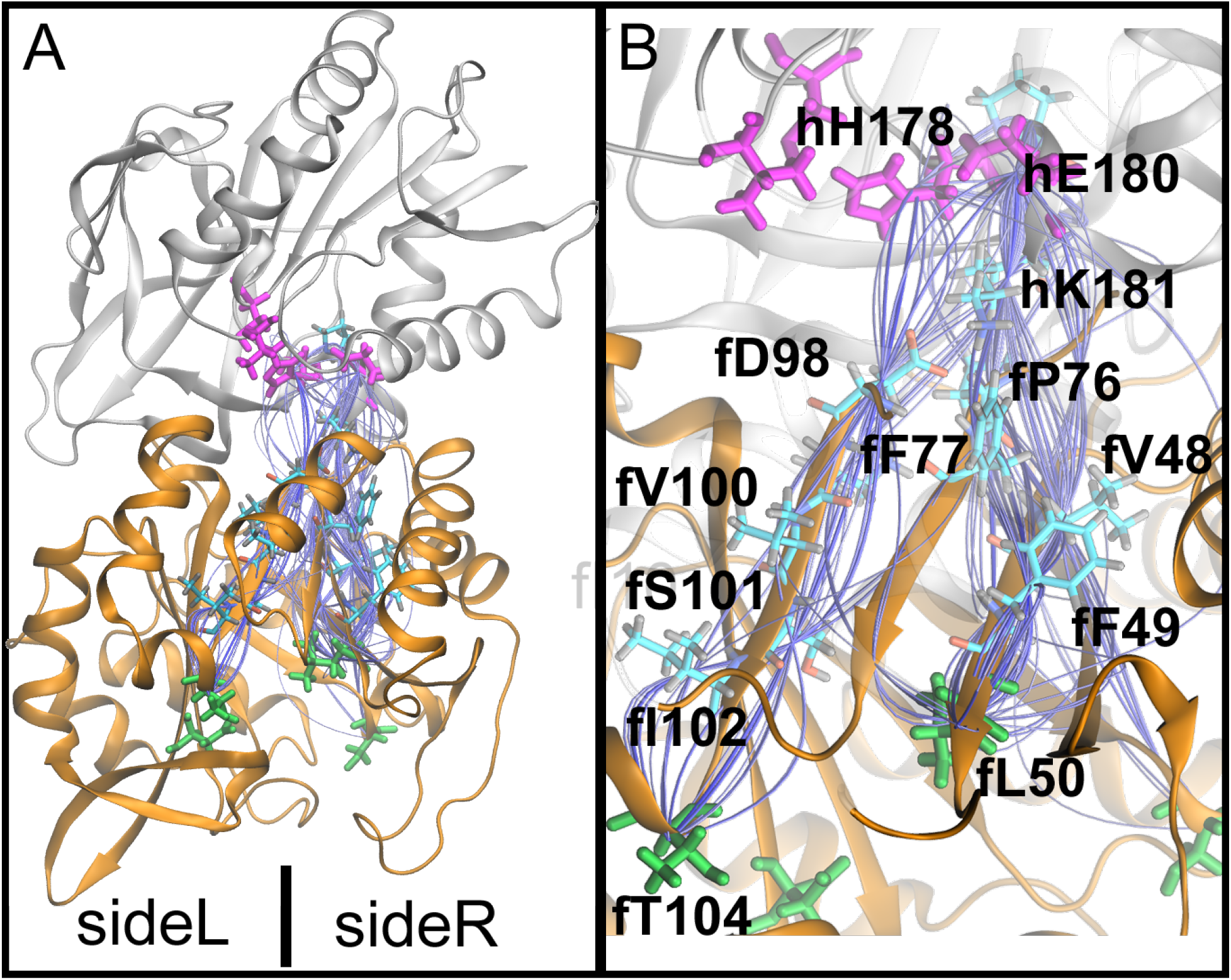
The 100 shortest direct allosteric paths determined based on the Hessian-based adjacency matric. A) A depiction of the full protein with paths dominantly sampling SideR of the protein. B) A close-up of the paths mainly propagating from fL50 and fT104 to hE180 and hH178.

### 3.3 Derivative Centrality Metric of the Hessian

Paths are an appealing approach to identify important residues between binding sites, but direct paths are not the only contributors to the covariance between source and sink residues. Using the Hessian as the adjacency matrix provides a unique analysis to compare to mutagenesis experiments that is unavailable when using other adjacency matrices. Given the physical interpretation of the finite Hessian elements, we can consider how the covariance between a source and sink is affected by changes in the Hessian elements, namely the spring constants between nodes. The details of this method, termed the derivative centrality metric, are provided in the Theoretical Framework section. This type of thinking parallels work from Rocks *et al.* but has not been applied directly to allostery in a protein. ^89,90^

The derivative edge metric identifies the formation of new edges across the HisF-HisH interfaces as being impactful on the covariance between sources and sinks. A structural representation of the edges with large magnitude derivative values (*δ*_*edge*_) from Equation 5 is provided in Figure 5A. The highest density of large magnitude (green) edges span the heterodimer interface, specifically at a region identified previously by Amaro *et al.* and Rivalta *et al.* to be modulated by PRFAR-binding. ^36,84^ An increased frequency of the reported “breathing motion” impacting this region can be interpreted as a strengthening of the spring constants that span the interface in the vicinity. Results from the derivative centrality metric suggest that the strengthening of these spring constants will change the covariance between source and sink residues, thus indicating an indirect allosteric effect on the glutaminase active site due to PRFAR binding. In the context of IGPS, the interfacial residues can be considered a communication bottleneck between HisH and HisF and thus altering the connectivity at the interface will have a large impact on the allostery between active sites.

**Figure 5:**
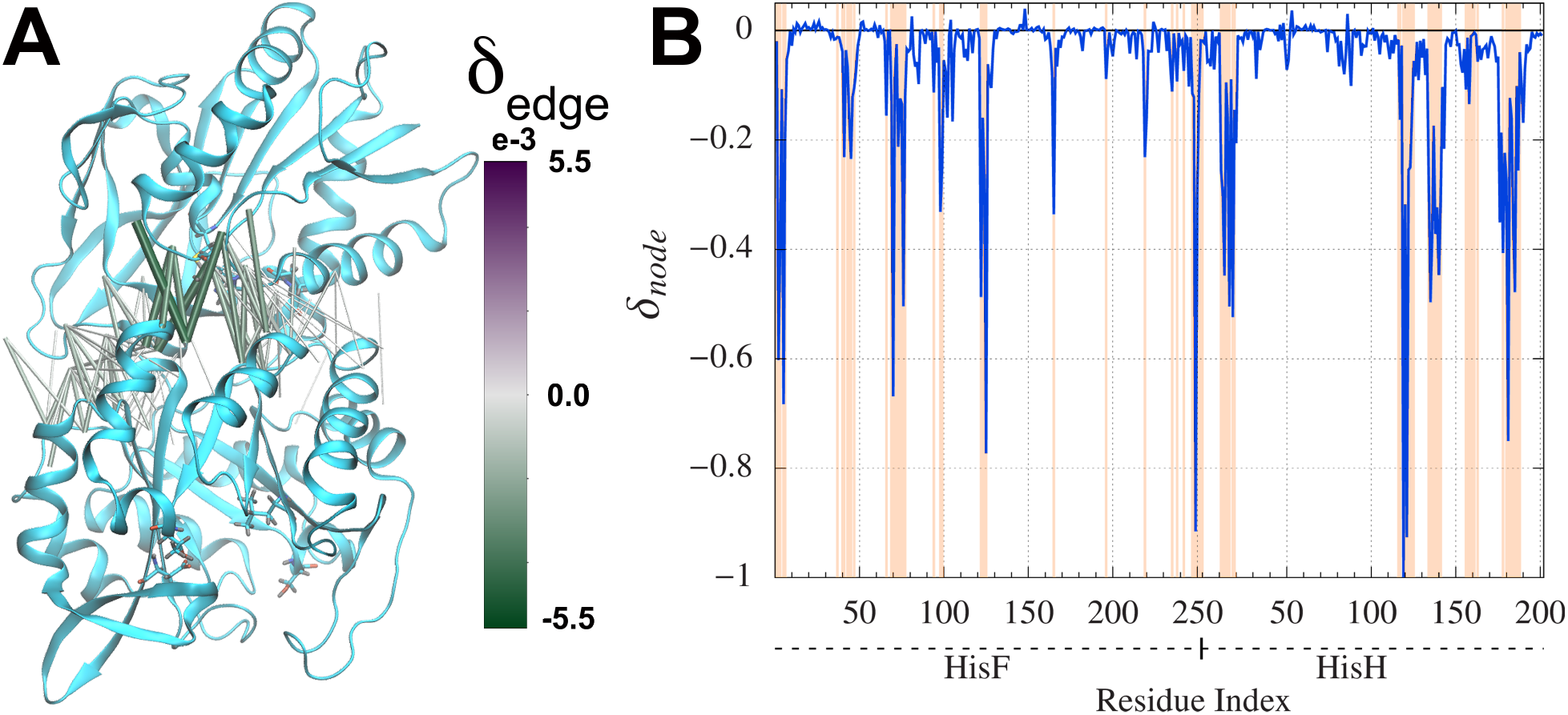
Hessian-based derivative centrality metric *δ*_*node*_ of IGPS apo. A) A structural representation edges with large derivative metric values B) Derivative node metric values as a function of residues. Residues that have connections across the HisF–HisH interface are colored by vertical orange lines.

The derivative node metric (*δ*_*node*_, Equation 6) for *apo* IGPS highlights the importance of a second position at the interface between HisF and HisH. These values are plotted as a function of residue number in Figure 5B with interfacial residues highlighted in orange. A cluster of three residues with large magnitude *δ*_*node*_ values are hP119 (*δ*_*node*_ = −1.00(4)), hM121 (*δ*_*node*_ = −0.9(1)) and fV125 (*δ*_*node*_ = −0.773(6)). These three residues sit at the interface between HisF and HisH, on sideL of the region discussed above. The interface is closer together at this position and the interfacial contacts in the Hessian matrix are observed to be stronger. The large magnitude *δ*_*node*_ values of these three residues suggest that the perturbation of their respective contacts will have a large impact on the covariance between sources and sinks. Due to the limited focus on this region of IGPS, we proffer this cluster of residues as potential targets for mutagenesis or inhibitory binding studies.

The path-based and derivative centrality metrics highlight different residues. This can be observed by noting that bridge residues, highlighted in orange, tend to be underrepresented in Figure 4B and highly represented in Figure 5B. The structural comparison also indicates that *P*_*node*_ highlights residues on sideR of the protein. On the other hand, the derivative centrality metric highlights only the interfacial residues, which span both sides of the protein. The correlation between these metrics is plotted in Figure S2. As presented in that figure, the majority of residues highlighted in the derivative metric are not observed in the direct paths between sources and sinks. The few exceptions to this are interfacial residues such as hK181 that are both at the interface, as well as in the direct paths. The frequency of such residues is relatively rare, thus we conclude that these two analyses provide different and complementary information. With this in mind, when considering how to disrupt or manipulate allostery, both direct and indirect effects should be understood: either the ensemble of allosteric paths can be directly disturbed by mutating a residue likely to be in the paths, or indirectly perturbed by changing the network contacts. Perturbations of areas considered to be important by the P_*node*_ and *δ*_*node*_ centrality metrics may be achieved through mutagenesis or inhibitory binding studies. It should be noted that these metrics account for the long-ranged correlations between binding sites. Therefore, they do not capture the effect of perturbing the direct, binding interactions between the protein and the effector molecule, although this is a considerable approach for disrupting allosteric regulation.

### 3.4 Comparison to Experimental Results for IGPS

The results from our simulations and centrality metrics on the effective harmonic coarsegrained Hessian compare favorably to mutagenesis experiments. To assess the mechanistic role of specific residues, we will focus on comparing our results to kinetic assay experiments of wild-type (WT) and mutant IGPS proteins. These assays compute the Michaelis-Menten enzymatic efficiency metric, k_*cat*_/K_*M*_, in the activated (effector bound) and basal states of the protein. The ratio of the enzymatic efficiency in these two states indicates the allosteric enhancement factor of the enzyme. The WT enzyme exhibits an allosteric enhancement factor of 4,500 to 4,900^31,73^ while most mutant proteins have a smaller enhancement factor relative to WT, indicating reduced allosteric activation of glutaminase activity by PRFAR (see Table 1).

**Table 1:** Residue importance for glutaminase related allostery in IGPS. A comprehensive list of published kinetic assay results for glutaminase activity of single point mutants of IGPS is included as well as our node centrality metrics, ***P***_*node*_ and *δ*_*node*_, for each of the mutated residues with error estimates in parentheses.

| Residue | Mutation | Ratio of Activated/Basal<br>$k_{cat}/K_M$ | $P_{node}$ | $\delta_{node}$ |
| --- | --- | --- | --- | --- |
| fR5 | fR5A<sup>a,b</sup> | 870 | 0.006(2) | -0.68(5) |
|  | fR5H<sup>a,b</sup> | 71 |  |  |
|  | fR5K<sup>a,b</sup> | 440 |  |  |
| fV12 | fV12A<sup>c</sup> | 3095 | 0.0132(10) | -0.004(4) |
| fK19 | fK19A<sup>c</sup> | 92 | 0.0000 | 0.004(2) |
|  | fK19A<sup>a,b</sup> | 45 |  |  |
|  | fK19R<sup>a,b</sup> | 110 |  |  |
| fV48 | fV48A<sup>c</sup> | 139 | 0.19(2) | -0.071(7) |
| fD98 | fD98A<sup>a,d</sup> | 2 | 0.16(6) | -0.33(7) |
|  | fD98A<sup>c</sup> | 54 |  |  |
| fK99 | fK99A<sup>a,b</sup> | 9100 | 0.08(2) | -0.25(2) |
|  | fK99R<sup>a,b</sup> | 700 |  |  |
| fT104 | fT104A<sup>a,e</sup> | 40 | 0.18(3) | 0.019(3) |
| fQ123 | fQ123A<sup>a,d</sup> | 2000 | 0.0000 | -0.16(6) |
| hN12 | hN12A<sup>a,d</sup> | 100 | 0.001(2) | -0.30(9) |
| hK181 | hK181A<sup>a,d</sup> | 2000 | 0.59(6) | -0.75(14) |
| WT<sup>b,d</sup> |  | 4500 – 4900 | – | – |

Mutant residues are reported using *T. maritima* IGPS notation. <sup>a</sup> denotes mutations performed in *S. cerevisiae* IGPS. <sup>b</sup> is from Ref. 73, <sup>c</sup> is from Ref. 31, <sup>d</sup> is from Ref. 77 and <sup>e</sup> is from Ref. 84.

The mutation of a residue found in an allosteric pathway is hypothesized to have an effect on the allosteric activation of the enzyme. Therefore, an adjacency matrix which corroborates experimental mutagenesis results holds value for the biological community. Of the residues studied by mutagenesis (Table 1), we find that only residues hK181, fT104, fV48, and fD98 have a significant (above 10%) probability of being in the direct paths between the PRFAR binding pocket and glutaminase active site. Of these, single point mutations of fT104, fV48 and fD98 to alanines diminish allosteric enhancement factor by over an order of magnitude compared to WT. This is consistent with the picture that altering residues in the allosteric paths found in the Hessian disrupts the ability of the enzyme to properly convey covariance between pockets. The Pearson and *r*_*MI*_ adjacency matrices struggle to corroborate these experimental results (see table S1).

Interestingly, a single point alanine mutation to hK181 only causes a factor of two decrease in allosteric enhancement factor when compared to WT. This result calls into question the importance of the salt-bridge between hK181 and fD98 that has been previously implicated as of extreme importance in IGPS allosteric paths. As was mentioned in the previous paths sections, we find that the residues fP76 and hK181 are more sampled in the paths than fD98. Additionally, in the IGPS *apo* Hessian, the fP76-hK181 and fD98-hK181 spring constants are 7.322 and 0.411 kcal mol^−1^Å^−2^, respectively. The interaction between fP76 and the aliphatic portion of hK181’s side chain is stronger than the salt bridge observed between fD98-hK181. We hypothesized that these trends in interactions would be well maintained in an hK181A mutant. To test this, we performed a simulation of the mutant and found a comparable force constant for the fP76-hA181 edge (2.844 kcal mol^−1^Å^−2^) and no force constant between fD98-hA181. Therefore, the alanine mutation of hK181 constitutes a small perturbation to the direct allosteric paths, which explains the small decrease in experimentally observed allosteric enhancement factor.

Residues that are at the HisF–HisH interface but not in allosteric paths can also have a large effect on the covariance between sources and sinks as demonstrated by our derivative metric. In Table 1, interfacial residues consist of fR5, fK99, fQ123 and hN12. The *δ*_*node*_ values for these residues are respectively −0.68(5), −0.25(2), −0.16(6), and −0.30(9), indicating that perturbations to the connectivity of these residues will have a significant impact on the observed allostery. Experimentally, mutations to these residues are all shown to have a large effect on allosteric enhancement relative to WT. The mutations have changed the interaction network around the mutated residue, which, in turn, caused a change to the covariance between PRFAR binding pocket residues and glutaminase active site residues. Interestingly, mutation of fK99 can either have a decrease (fK99R) or an increase (fK99A) in activity relative to WT. The derivative metric does not indicate how the change in covariance affects allosteric enhancement just that it will change. It will thus be of interest to study fK99 further to investigate how these two mutants affect the covariance.

If the path and derivative centrality metrics provide a complete picture of allostery in IGPS, then residues not in paths and not at the HisF–HisH interface will have little effect on the experimentally measured allosteric enhancement factor. Residue fV12 is observed in only 1% of paths and has a *δ*_*node*_ of only −0.04(4). A valine to alanine mutation at this position has little effect on the allosteric enhancement factor relative to WT, suggesting that the *P*_*node*_ and *δ*_*node*_ results do provide a complete picture for this residue. In contrast, fK19 has *P*_*node*_ = 0 and *δ*_*node*_ = 0.04(2) yet all three mutations listed in Table 1 have a large effect on allosteric enhancement factor. Amaro *et al.* performed aaMD simulations to model the unbinding of the PRFAR ligand, from which fK19 was observed to play an important role in the recognition and binding of the effector molecule. ^84^ A mutation to the fK19 residue is likely to decrease the observed allosteric enhancement factor by simply reducing HisF’s binding affinity for PRFAR. This effect would not be detectable with our metrics because it does not represent a perturbation to the allosteric, long-range correlations between binding pockets.

## Conclusions

In this work, we provided evidence that the effective coarse-grained Hessian is the appropriate graph Laplacian to consider in the context of allostery. The Hessian only contains finite values for short-ranged physical interactions and can be rigorously tied to the covariance for harmonic systems. We use a previously developed coarse-graining protocol, hENM, to compute the best effective harmonic Hessian that captures the residue-level covariance of all-atom molecular dynamics systems.

With the Hessian as the graph Laplacian, we develop two centrality metrics to highlight important residues that contribute to allostery. Both of these metrics are applied to the IGPS protein to investigate interactions between the PRFAR binding pocket and the glutaminase active site. The first of these metrics is based on direct path sampling and recapitulates the known importance of sideR in the allostery of IGPS. Paths in the Hessian-based adjacency matrix are found to be significantly longer and to be less diluted than paths found on two previously used adjacency matrices.

The second centrality metric we develop is based on the derivative of the covariance as a function of a given Hessian element. This metric is motivated by mutagenesis experiments in which one perturbs the interactions around a mutation site as compared to wildtype. This metric identifies residues at the interface between HisF and HisH as being important for the allosteric network between the PRFAR binding pocket and the glutaminase active site. The correlation between the derivative and path centrality metrics is found to be minimal, suggesting that these two metrics provide different yet important information about the covariance between the two pockets.

Results from the path and derivative centrality metrics on the effective Hessian corroborate and functionally explain previous mutagenesis experiments. Interestingly, the combination of experimental and simulation results suggest that fP76-hK181 is a more important HisF– HisH interfacial connection for allosteric paths than the previously implicated fD98-hK181 salt bridge. Additionally, we propose a novel region for targeted mutagenesis or inhibitory binding studies based on the results obtained from our two centrality metrics.

It is well agreed upon that allosteric networks exist regardless of the bound state of the protein, therefore this work focused on the apo IGPS state. Further exploration of how the Hessian network is altered in the holo IGPS state awaits. Additionally, the authors are interested in studying ways in which metastable conformational states may alter Hessian network preferences and provide insight into conformational shifts that are more suitable for the allosteric network.

## Acknowledgement

This work used the Extreme Science and Engineering Discovery Environment (XSEDE), which is supported by National Science Foundation grant number ACI-1548562. We would specifically like to acknowledge the San Diego Supercomputer Center Comet and Pittsburgh Super Computing Bridges resources used under XSEDE allocation CHE160008 awarded to MM.

## Supporting Information Available

Additional information is provided including: details of the computational methods employed to perform all simulations, further details of the hENM procedure, further details on the allosteric path sampling algorithm, figure demonstrating the agreement between covariance of aaMD and CG models and figure demonstrating lack of correlation between the two centrality metrics discussed in this paper. All code to perform these analyses is provided at: https://github.com/mccullaghlab/Allostery-Analysis.

