## Supporting Information for "Residue-Level Allostery Propagates Through the Effective Coarse-Grained Hessian"

#### Computational Methods

##### Heterogeneous elastic network models

Here we will closely follow the procedure known as heterogenous ENM (hENM) to generate an effective harmonic Hessian that best reproduces the aaMD measured covariance.[1] Our adapted hENM algorithm is as follows:

1. Initiate a trial $N \times N$ force constant matrix, $\boldsymbol{k}$.
2. Create the $3N \times 3N$ Hessian matrix, $\boldsymbol{H}$ using the tensor matrix elements $\boldsymbol{H}_{ij} = -k_{ij} \cdot \hat{r}_{ij} \otimes \hat{r}_{ij}$ for $i \neq j$ and $\boldsymbol{H}_{ii} = \sum_{j \neq i} \boldsymbol{H}_{ij}$

3. Get the predicted $3N \times 3N$ covariance, $\boldsymbol{C}$, by performing the Moore-Penrose psuedo inverse of $\boldsymbol{H}$, $\boldsymbol{C} = T\boldsymbol{H}^{+}$.

4. Update the force constants based on

$$k_{ij}^{n+1} = k_{ij}^{n} - \alpha \left( \frac{1}{\sigma_{ij}^{n}} - \frac{1}{\sigma_{ij}^{target}} \right) \tag{SI1}$$

where $\sigma_{ij}^{n} = \hat{r}_{ij} \left[ \boldsymbol{C}_{ii}^{n} + \boldsymbol{C}_{jj}^{n} - \boldsymbol{C}_{ij}^{n} - \boldsymbol{C}_{ji}^{n} \right] \hat{r}_{ij}$ is the variance of node $i$ and $j$ projected along the separation vector and $\alpha$ is a minimization step size parameter (we found $\alpha = 1 \times 10^{-2}$ to yield stable convergence).

5. Repeat steps 2-4 until a converged Hessian matrix has been produced.

#### All-Atom MD Simulations

All-atom, explicit solvent MD simulations are performed for the *apo* substrate state of *T. maritima* IGPS (PDB: 1GPW).[2] Specifically, chains C and D are used from this crystal structure. Crystallographic waters are maintained. The active site mutation, fN11D, present in this crystal structure was converted back to wild type. Additionally, MD simulations were performed of an artificial hK181A mutant. Starting from the 1GPW structure, the mutation was performed by removing sidechain atoms associated with the lysine and allowing tleap to replace the missing atoms with alanine sidechain. This simulation followed the same procedure as *apo*, detailed below.

The simulations are performed using the GPU-enabled AMBER18 software[3] and the ff14SB[4] parameters for proteins atoms. Using tleap, the starting structure is solvated in a TIP3P water box with at least a 12 Å buffer between the protein and periodic images. Sodium and chloride ions are added to neutralize charge and maintain a 0.10 M ionic concentration. Direct nonbonding interactions are calculated up to a 12 Å distance cutoff. The SHAKE algorithm is used to constrain covalent bonds that include hydrogen.[5] The particle-mesh Ewald method[6] is used to account for long-ranged electrostatic interactions. Before simulation began, a two stage minimization was performed: (1) 10,000 steps of conjugate gradient optimization were performed to minimize water positioning (substrate atoms restrained with a 75 kcal $mol^{-1}$ $Å^{-2}$ force constant) and (2) an additional 10,000 minimization steps with no restraints applied. The system was slowly heated from 25 K to 303 K over 1 ns. Additionally, 4 ns of NVT simulation was performed to equilibrate the cubic box volume. Finally, the simulations were run in the NTP ensemble, using the Langevin dynamics thermostat and Monte Carlo barostat to maintain the systems at 303 K and 1 bar. A 2 fs integration time step is used, with energies and positions written every 5 ps. An initial 500 ns simulation was run, from which the structure at 250 ns was used to initialize three more independent trajectories. These new trajectories were given different random number seeds and were run for an additional 250 ns. We have a total of 1 $\mu$s of trajectory to analyze the *apo* state of IGPS as well as the hK181A mutant system.

### Theory

#### Linear Response Solution to Allosteric Pathways

#### Path Sampling

The emphasis of the ensemble of paths makes sampling the paths an apt way of studying them. For this purpose, we present a Markov chain simulation to sample the paths. This work is in contrast to previous studies, namely WISP,[7] that focus on finding the exact shortest paths up to some number. Instead, we statistically sample a distribution of paths. This algorithm is shown to be more efficient than using WISP to accomplish the same result.

The set of objects that are studied using the Markov chain simulation is the set of paths from a predefined source and sink nodes. Each path is then assigned an effective length. For the purpose of this work we consider some adjacency matrix $\boldsymbol{A}$ and functionalize it into a cost matrix $\boldsymbol{\chi}$ in an analogous way as the covariance is related to the Pearson correlation. Namely,

$$\chi_{ij} = -\log\left(\frac{|A_{ij}|}{\sqrt{A_{ii}A_{jj}}}\right) \tag{SI2}$$

In this study, we consider the paths of three adjacency matrices: the covariance, $r_{MI}$, and the Hessian. The length $\ell$ of a path is defined as a sum over values in the cost matrix corresponding to the edges in the path. The distribution of the paths that are studied is of the form

$$P_{path} \propto \exp\left[-\ell/\tau\right], \tag{SI3}$$

where the free parameter $\tau$ is an effective temperature of the simulation. The Markov chain simulation we present is not limited to these definitions of path lengths and path probabilities; more complex functions of the weights and the path distributions can readily be used.

A trial move in the Monte Carlo simulation consists of randomly choosing a node in the graph that is neither the source nor sink node from a uniform distribution, and add/remove the node to/from the path if it is currently/not in the present path. Any trial move that destroys the path, as will happen when an edge with a zero in the adjacency matrix is used, the move is rejected. The following values associated with the probability of adding the selected node $n$ between each pair of connected nodes $i$ and $j$ in the path not containing $n$ is then calculated

$$p_{ij} = \exp\left[-\frac{\chi_{in} + \chi_{nj} - \chi_{ij}}{\tau}\right]. \tag{SI4}$$

If the node is not in the current path, the trial move is a path with node $n$ randomly inserted between two connected nodes with the probability proportional to $p_{ij}$. Detailed balance is achieved by using a Metropolis algorithm whereby the trial path is accepted with a probability $P_{Met}$ of

$$P_{Met} = \min\left[1, \left(\sum_{\langle ij\rangle} p_{ij}\right)^{\pm 1}\right], \tag{SI5}$$

where the upper sign is for when the trial move removes a node from the path and the lower sign is for node addition.

The difference between using the Hessian as the adjacency matrix and the equation we derived relating the Hessian to the covaraince in terms of paths can be mocked in the sampling on a few levels. As a first pass, the Hessian can be treated as a $3N \times 3N$ object where an edge in a graph is described by a $3 \times 3$ submatrix. The nature of the Hamiltonian we are using to find spring constants of the system produces a $3 \times 3$ submatrix describing the interaction of node $i$ and $j$ of the form $k_{ij}\hat{\boldsymbol{r}}_{ij}\hat{\boldsymbol{r}}_{ij}^{\intercal}$. The weight $w$ of a path can then be considered to be the product of these spring constants, treated in the same fashion as above, as

$$\begin{aligned} w &= \prod_{\langle i,j \rangle} \frac{k_{ij}}{\sqrt{k_{ii}k_{jj}}} \hat{\boldsymbol{r}}_{ij}\hat{\boldsymbol{r}}_{ij}^{\intercal} \\ &= \hat{\boldsymbol{r}}_{i_1 i_2}\hat{\boldsymbol{r}}_{i_{M-1} i_M}^{\intercal} \prod_{\langle i,j,k \rangle} \cos\theta_{ijk} \prod_{\langle i,j \rangle} \frac{k_{ij}}{\sqrt{k_{ii}k_{jj}}}, \end{aligned} \tag{SI6}$$

where $\langle i,j,k \rangle$ is the set of all three consecutive nodes in the path. The two products in this equation can be interpreted as the scalar weight of the path and the tensor part is treated separately. Taking the length of the path to be equal to $\ell = -\log w$, these paths can similarly sampled with the following change to the $p_{ij}$ defined above,

$$p_{ij} = \left( \frac{\cos\theta_{i-1,i,n} \cos\theta_{i,n,j} \cos\theta_{n,j,j+1}}{\cos\theta_{i-1,i,j} \cos\theta_{i,j,j-1}} \right)^{1/\tau} \exp\left[ -\frac{\chi_{in} + \chi_{nj} - \chi_{ij}}{\tau} \right]. \tag{SI7}$$

Notice that this definition of path length is not trivially studied by exhaustive search algorithms such as that proposed by WISP, but it readily studied using this Markov chain simulation framework.

A simulation of the paths consists of $N$ trial moves between writing the path to file, where $N$ is the number of nodes in the graph, and then $10^6$ paths are generated to sample the ensemble. The choice of $\tau$ is dependent on the desired ensemble of paths to sample. For the present study, $\tau$ is chosen such that the first thousand shortest paths are well sampled so that our results can be compared to that of WISP.

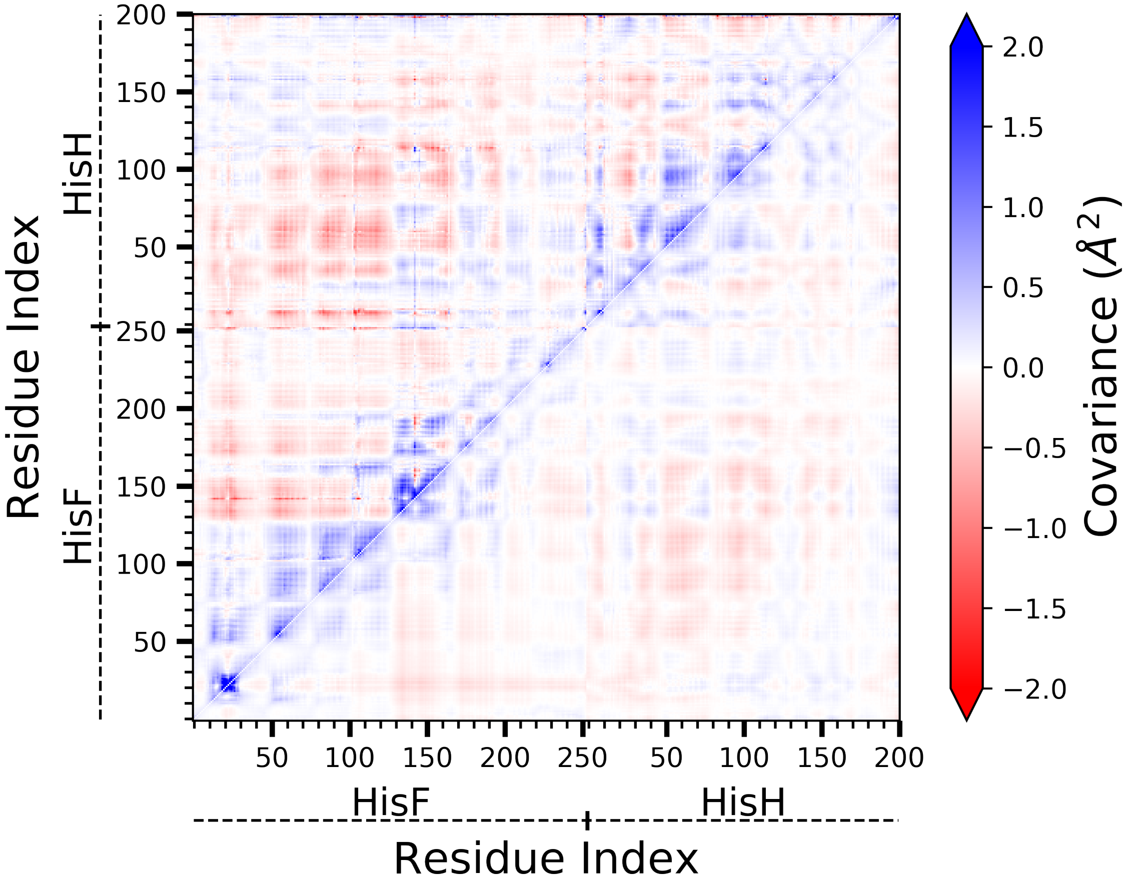


Figure S1: Covariance matrices computed through the hENM procedure (bottom right) and raw simulation (top left) from 1 $\mu$s all-atom molecular dynamics of *apo* IGPS.

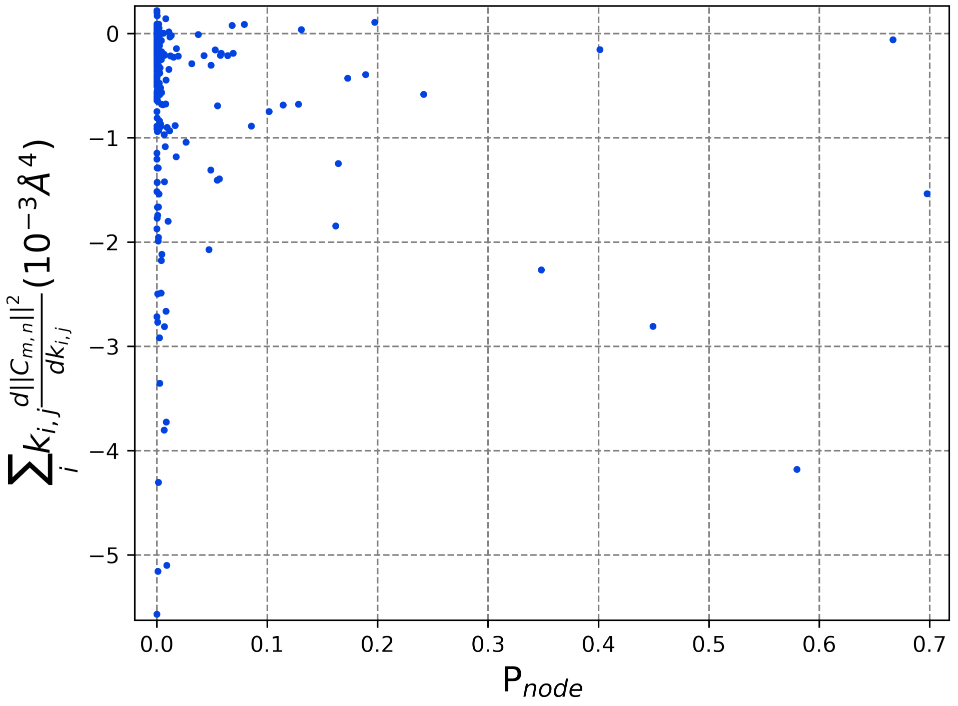


Figure S2: Correlation of connectivity matrices $P_{node}$.

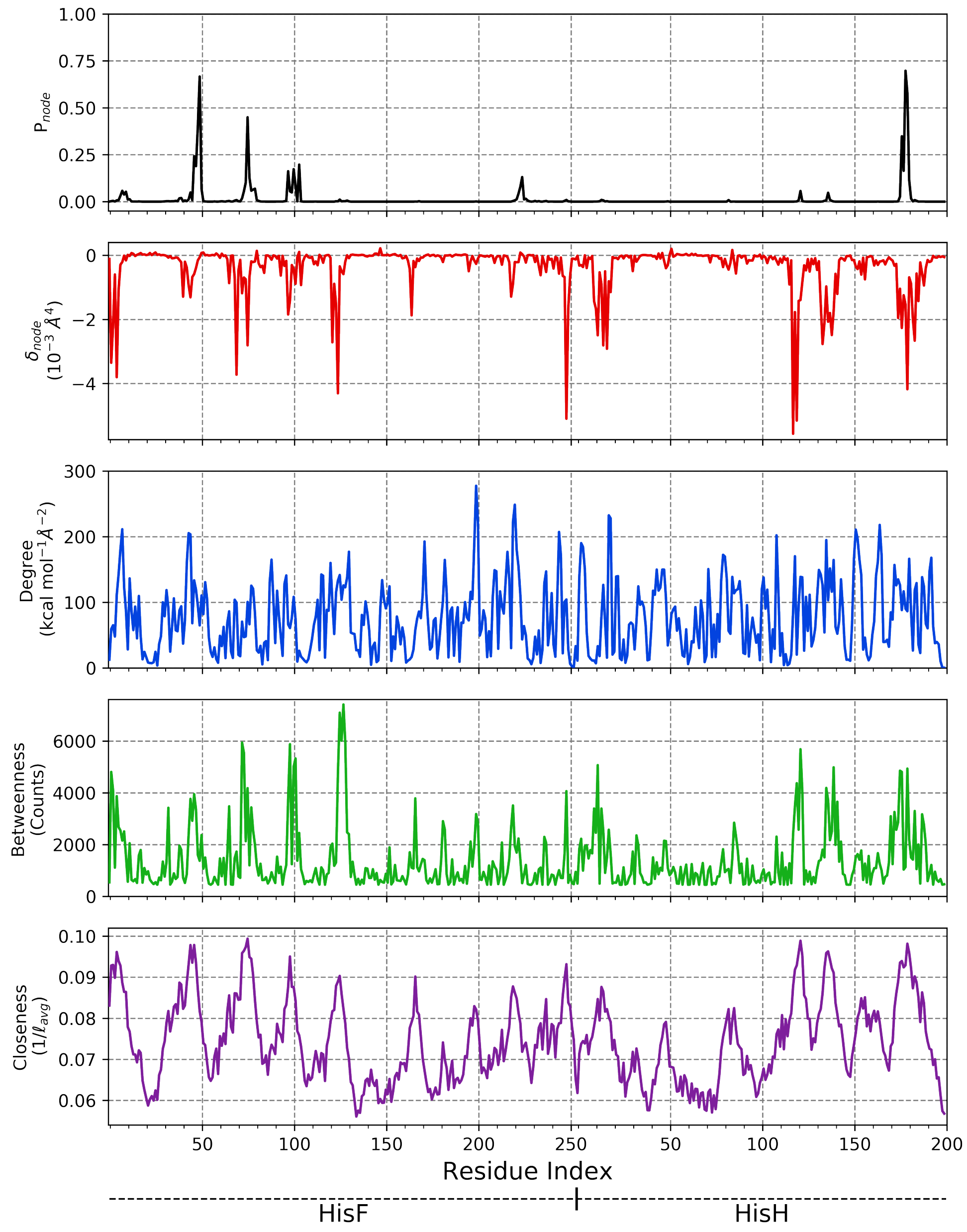


Figure S3: Comparison of $P_{node}$ and $\delta_{node}$ centrality metrics (reported in the article, top two plots) to commonly used centralities (Degree, Betweenness, and Closeness, bottom three plots). All analyses were performed using the *apo* IGPS Hessian adjacency matrix. These common metrics do not directly account for allostery and, so, residues with large magnitude values are not gua directly important

Table S1: Residue importance for glutaminase related allostery in IGPS. Comparison of $P_{node}$ centralities resulting from different connectivity matrices for selected mutated residues.

| Residue | Mutation | Activated/Basal $k_{cat}/K_M$ | $P_{node,Hessian}$ | $P_{node,r_{MI}}$ | $P_{node,Pearson}$ |
|---|---|---|---|---|---|
| fR5 | fR5A$^{a,b}$ | 870 | 0.0068 | 0.0000 | 0.0000 |
| | fR5H$^{a,b}$ | 71 | | | |
| | fR5K$^{a,b}$ | 440 | | | |
| fV12 | fV12A$^{c}$ | 3095 | 0.0131 | 0.0001 | 0.0000 |
| fK19 | fK19A$^{c}$ | 92 | 0.0000 | 0.0000 | 0.0000 |
| | fK19A$^{a,b}$ | 45 | | | |
| | fK19R$^{a,b}$ | 110 | | | |
| fV48 | fV48A$^{c}$ | 139 | 0.1891 | 0.0000 | 0.0000 |
| fD98 | fD98A$^{a,d}$ | 2 | 0.1620 | 0.0000 | 0.0000 |
| | fD98A$^{c}$ | 54 | | | |
| fK99 | fK99A$^{a,b}$ | 9100 | 0.0547 | 0.0000 | 0.0000 |
| | fK99R$^{a,b}$ | 700 | | | |
| fT104 | fT104A$^{a,e}$ | 40 | 0.1973 | 0.9590 | 0.9819 |
| fQ123 | fQ123A$^{a,d}$ | 2000 | 0.0000 | 0.0000 | 0.0000 |
| hN12 | hN12A$^{a,d}$ | 100 | 0.0004 | 0.0000 | 0.0000 |
| hK181 | hK181A$^{a,d}$ | 2000 | 0.5797 | 0.0000 | 0.0000 |

Mutant residues are reported using *T. maritima* IGPS notation. [a] denotes mutations performed in *S. cerevisiae* IGPS. [b] is from Ref. 8, [c] is from Ref. 9, [d] is from Ref. 10 and [e] is from Ref. 11.
